# Exploration of the antimicrobial synergy between selected natural substances on *Streptococcus mutans* to identify candidates for the control of dental caries

**DOI:** 10.1101/2021.11.19.469353

**Authors:** Alisha Prince, Soumya Roy, David McDonald

## Abstract

Dental caries is caused by dental plaque, a community of micro-organisms embedded in an extracellular polymer matrix as a biofilm on the tooth surface. Natural products that are widely available could be used as an alternative or adjunctive anti-caries therapy. Sometimes, when two products are used together, they yield a more powerful antimicrobial effect than the anticipated additive effect. These synergistic combinations are often better treatment options because individual agents may not have sufficient antimicrobial action to be effective when used alone. Cranberries contain phenolic compounds like proanthocyanidins (PAC) that disrupt biofilm formation. Manuka honey has high concentrations of the agent methylglyoxal, which is cariostatic. Because these agents have varied modes of antimicrobial action, they show potential for possible synergistic effects when paired.

Various cranberry extracts were tested pairwise with manuka honey or methylglyoxal by well-diffusion assays and 96-well checkerboard assays in the presence of *Streptococcus mutans* to test for synergy. Synergy was demonstrated in two of the cranberry extracts paired with manuka honey. The synergistic combinations found in this research thus can be considered as candidates for the formulation of a dentifrice that could be used to inhibit the formation of dental plaque and thereby avoid the development of caries.

**Significance and Impact:** The emergence of bacteria resistant to antimicrobial agents has led to a shortage of options when choosing effective treatment agents. Further, some antibiotics used at therapeutic doses can produce undesired side effects. However, by utilizing synergistic combinations of antimicrobial agents, the desired antimicrobial effects can often be produced without subjecting the patient to toxic or irritating doses of individual agents. *Streptococcus mutans* growth and biofilm formation is a major contributor towards dental caries. In this study, a synergistic combination of Manuka honey and cranberry extracts gives evidence that it can be used as an alternative or adjunctive anti-caries therapy.

## Introduction

As early as 1924, it was discovered that *Streptococcus mutans* forms a major part of dental plaque (1). *In vivo* studies with germ-free rats corroborated these findings, proving that tooth decay and gum disease are caused by bacteria (2). *S. mutans* is a facultative anaerobic gram-positive coccus. The virulence factors that contribute to their cariogenicity are (i) adhesion to tooth surfaces (ii) acidogenicity through glycolysis and (iii) acid tolerance (3). *S. mutans* have glucosyltransferases that can convert the dietary starches to extracellular polymers like glucan that can bind to the dental pellicle, which then allows for the attachment of other bacteria like *Streptococcus sanguis, Streptococcus oralis*, and *Lactobacillus sp*. (4). The fermentation of simple sugars by this bacterial consortium produces organic acids that can with time erode the enamel surface of the tooth and cause dental caries.

Many studies have been performed to test the activity of natural products like green tea, cacao bean, cranberry, cinnamon, garlic, honey, etc. against *S. mutants* (5–10). These can then be used as an alternative or adjunctive anti-caries agents.

Sometimes, when two products are used together, they yield a more powerful antimicrobial effect than the anticipated additive effect. These synergistic combinations are often better treatment options because individual agents may not have sufficient antimicrobial action to be effective when used alone or may require sufficiently high concentrations that they become toxic or irritating (11). Furthermore, exposure to dual agents greatly reduces the likelihood of the development of resistance by the target bacteria. Finally, it is also important to note that some natural substances, for example in this study cranberry and honey, have pleasant flavor profiles, which would likely lead to increased patient treatment compliance of a dentifrice substance containing them.

## Materials and Methods

### Materials

All cranberry extracts were provided by Ocean Spray Cranberries Inc. The cranberry extracts were labelled by the manufacturers as “SWP”(subcritical water extract of presscake), “SWF” (subcritical water extract of fruit), “SWPE” (subcritical water extract of presscake with tannase), “Type R” (resin extract) and “RE” (resin extract with tannase). The extracts vary based on their composition and methods of extraction (12). For the purpose of this experiment, we purchased manuka honey (Manuka Health, New Zealand) containing 550+ ppm methylglyoxal (MGO). Methylglyoxal (Sigma, USA) was added to distilled water to prepare a 550 parts per million solution of methylglyoxal, equivalent to the label-indicated content in the manuka honey. Cinnamon extracts labelled “Cinn-HA” and “Cinn Oil” (McCormick & Company, USA), neem seed oil (NeemAura, USA) as well as commercially available mouthwashes Colgate Total, Listerine and ACT mouthwashes were also used. To prepare a mix of extracts, 750μl of one extract was added to 750μl of another extract in a sterile Eppendorf tube and mixed well. Cranberry, cinnamon, manuka honey, MGO, neem extracts were used in combinations and each combination was tested in separate assays. Distilled water was used as a negative control. Chlorhexidine (2% v/v chlorhexidine gluconate) was used as a positive control.

Brain Heart Infusion (BHI) agar plates were used to culture *S. mutans* (ATCC 25175) by placing streak plates in a 5 % carbon dioxide (CO_2_) incubator at 37°C overnight. To standardize the bacterial suspension used in each assay, a 0.5 McFarland standard was prepared (13).

### Agar Well Diffusion Assay

100μl of *S. mutans* at a turbidity of 0.5 McFarland standard were added to each BHI agar test tube and poured into 20 petri plates. The plates were allowed to sit at room temperature for 2 hours. Using a sterile cork-borer, five wells were punched in the BHI agar of every petri dish. A template was used to ensure that the wells were equidistant from each other. In each petri dish, 95μl of cranberry, manuka honey, cranberry-manuka honey mixture, water and chlorhexidine were pipetted into the wells (Figure 1). After allowing them to diffuse for 30 minutes, they were placed in a CO_2_ incubator at 37°C overnight. This method was repeated with neem, cinnamon, MGO and the other cranberry extracts. To compare the efficacy of the most antimicrobial combination with commercial mouthwash, 95μl of the cranberry and manuka honey mixture was pipetted in one well and 95μl of Colgate Total™, Listerine™ and ACT™ mouthwashes were pipetted into the three other outer wells respectively. After a 24hr incubation period, the plates were photographed individually with a ruler, which served as a reference while measuring zones of inhibition. Image analysis was done using Adobe Photoshop CS6® (Adobe Systems) and Image J (14). The borders around the zones of inhibition weren’t always well defined. A photo training process similar to that described by Jorgensen et al (15) was used to read pixel values and demarcate the zones of inhibition. In analyzing the data, the goal was to detect variations in zone diameters between the extracts used separately and the extracts used in combination. Analysis of Variance (ANOVA) was used to detect if the differences were statistically significant. “Petri Dish ID” was added as a block factor as there could be some variances between each petri dish. This made the study a two factor ANOVA. “Blocks” and “substance added” were the independent variables and “diameter of zone of inhibition” was the dependent variable. This study is a Mixed Model ANOVA because “petri dish ID” is a random effect.

**Fig. 1.**
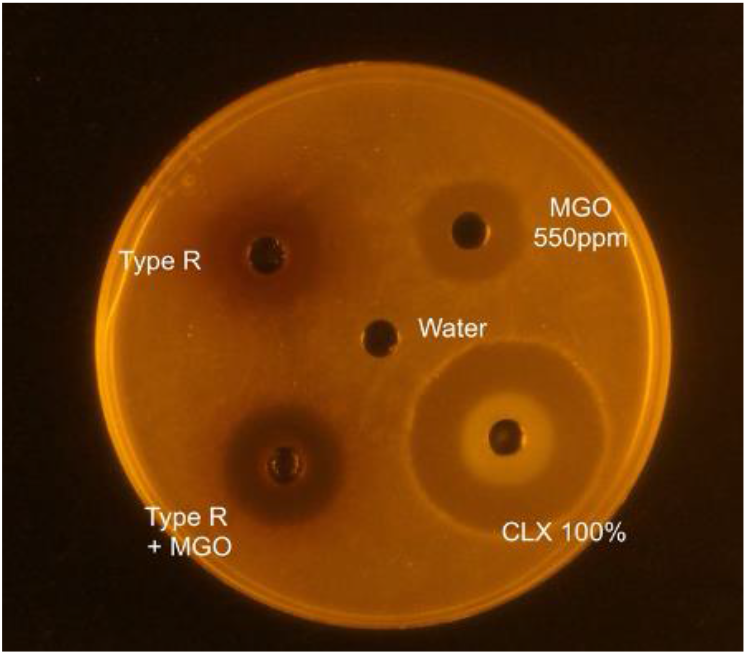
Well diffusion assay: cranberry Type R and MGO

In RStudio (16), using the lme4, lmerTest and lsmeans packages, Type III Analysis of Variance with Satterthwaite’s method was performed. This was followed by a post-hoc Tukey test to compare diameters of the zones of inhibition and detect synergy with a confidence interval of 95%.

The well diffusion and checkerboard assays with each combination was completed at least three times to make the data more robust and to assure the assay’s repeatability.

### Checkerboard Assay

Serial dilutions of bioactive agents were prepared with 600μl of sterile water. Corning Falcon™ polystyrene 96 well microplates with nontreated surfaces (Fisher Scientific, USA) were used in a clean laminar flow hood for the checkerboard assay to check for synergy as described by (17). 100μl of BHI broth were added to each well of the 96-well plate. 20 μl of *S. mutans* at 0.5 McFarland standard turbidity were added to each well, except the last column (column 12) of the well. These wells served as the negative growth control. Column 11 containing only media and the bacteria served as a positive growth control. Of each concentration of cranberry extract 50μl was added to wells in separate columns (columns 1 to 8). 50μl of MGO concentrations were added to rows A to H. This formed a checkerboard pattern with varying concentrations of cranberry in the columns, combined with varying concentrations of MGO in the rows (Figure 2). This 96-well plate was then placed overnight in a 5% (v/v) carbon dioxide (CO_2_) incubator at 37°C. After 18 to 24 hours, the 96-well plate was analyzed for bacterial growth. The first wells without any growth in each row and column, adjacent to the wells with bacterial growth form the “growth/no growth interface.” The first concentration of a bioactive agent without any bacterial growth is the Minimum Inhibitory Concentration (MIC). If A and B refer to the individual agents, the fractional inhibitory concentration (FIC) index was calculated for each well along the growth/no growth interface using the formula:

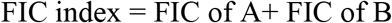

where,

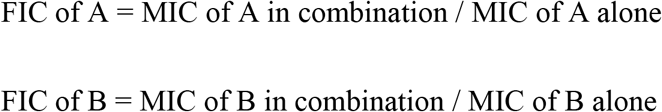

**Fig. 2.**
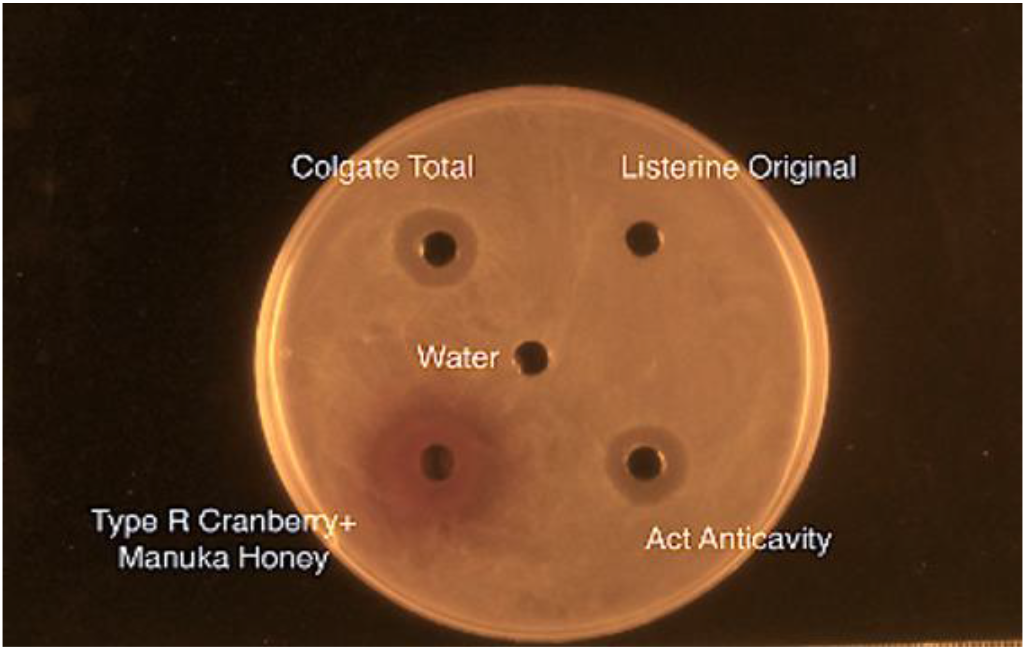
Well diffusion assay with cranberry Type R and Manuka honey combination, and commercial mouthwashes

Three methods were used to interpret the checkerboard assay and determine synergy. (i) Mean FIC index (18) (ii) Lowest FIC Index (19) (iii) Two Well Method (20). This methodology was applied to six different cranberry extracts, two cinnamon extracts, manuka honey, MGO and neem and studied in combinations.

## Results

Of the six cranberry extracts tested, two extracts when paired with manuka honey, showed stronger antimicrobial action in the well-diffusion assay. The zones of inhibition around the cranberry-manuka honey combination wells were larger than those around the cranberry and manuka honey wells, respectively. This difference was statistically significant (p < 0.0001) and repeatable, with consistent results when the experiments were performed in triplicate.

No synergy was found with neem and cinnamon extracts, where the zones of inhibition of their combinations with manuka honey were significantly smaller than the extracts used alone.

The cranberry extract – manuka honey combination was subjected to a well diffusion assay along with commercially-available mouthwash. The zones of inhibition around the cranberry-manuka honey combination was much larger than those around the mouthwashes (Figure 3). The cranberry and manuka honey combination (***) had statistically larger (p<0.0001) zones of inhibition than the mouthwashes (Figure 4). Colgate Total and ACT mouthwashes had statistically similar (p 0.2927) zones of inhibition (**). Listerine had a smallest (p<0.0001) zone of inhibition (*) among the mouthwashes tested in a well diffusion assay.

**Fig. 3.**
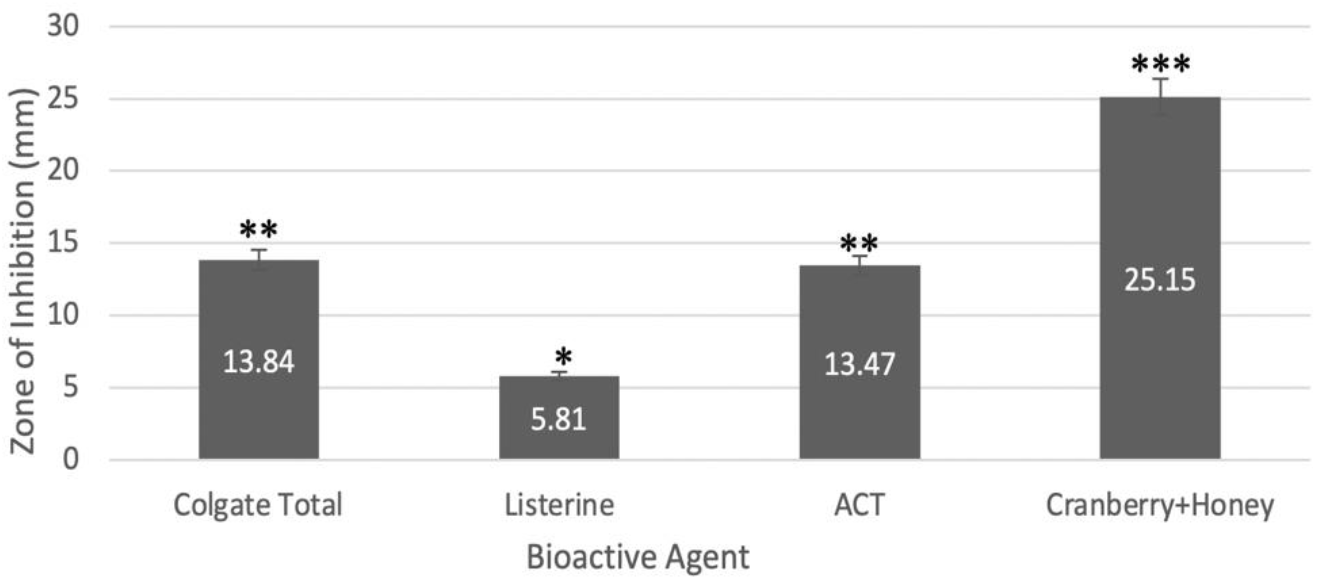
Zones of inhibition around wells containing cranberry and Manuka honey combination, and commercial mouthwashes in a well diffusion assay with *S. mutans*

**Fig. 4.**
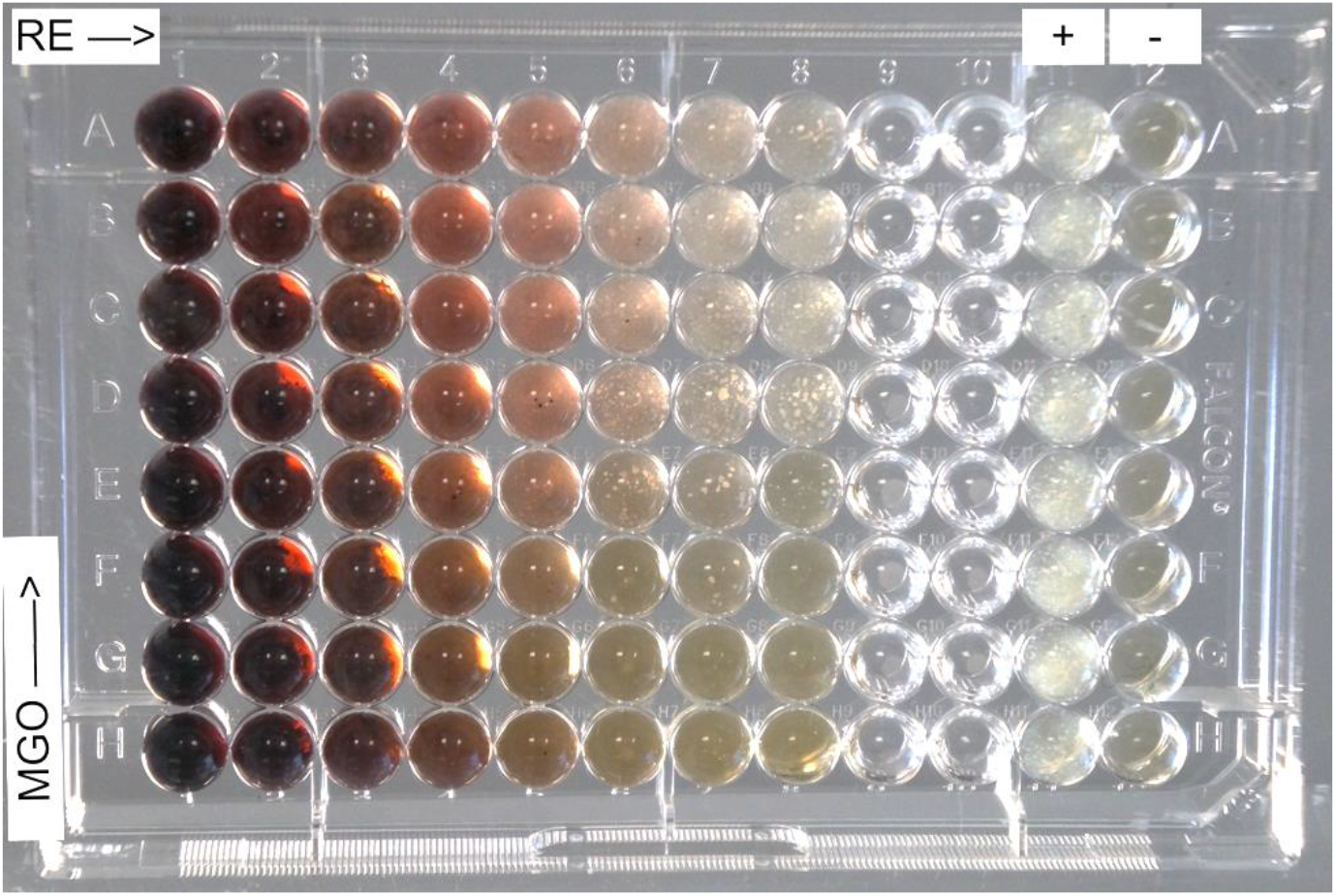
Checkerboard Assay with Cranberry serial dilutions in columns and MGO serial dilutions in rows

Cranberry extracts, neem and cinnamon extracts were respectively paired with MGO (the bioactive agent in manuka honey) in well diffusion assays. While neem and cinnamon did not show significant synergy, two of the cranberry extracts showed significantly (p<0.0001) larger zones of inhibition when used in combination with MGO, than when used alone.

The extracts that showed antimicrobial synergy were then subjected to a 96-well plate checkerboard assay. Three cranberry extracts when combined with MGO, showed synergy in all three methods of assessment. The mean FIC index for cranberry extracts combined with MGO were 0.6, 0.5 and 0.8 respectively, which were less than 1, indicating synergy (21). The lowest FIC indices were 0.38, 0.27 and 0.5, which were less than or equal to 0.5, indicative of synergy. This study was replicated and showed similar data. The MIC of MGO was 0.5 of its original 550ppm concentration. Two cranberry extracts had MICs of 1 (their original concentration) and the third cranberry extract had an MIC of 0.5. For a cranberry extract with an MIC of 1, serial dilutions of intermediate concentrations (0.75, 0.38, 0.19, etc.) were added to the checkerboard assay. This study gave an MIC of 0.75 for cranberry and 0.38 for MGO, indicating that for cranberry extract 0.75< MIC <1 and for MGO 0.50< MIC <1. The mean FIC index for this study remained at 0.5 and the lowest FIC index was 0.28.

## Discussion

Cranberry extracts R and RE when paired with Manuka honey and MGO respectively showed synergy in both the well-diffusion assay and the checkerboard assay. This may be the consequence of their varied modes of action being complementary against *S. mutans*. Cinnamon and neem did not show synergy in these assays. But they showed antimicrobial activity when used alone. Perhaps future studies will reveal how they can be used in combination with other antimicrobial agents to achieve a level of antimicrobial activity useful for dental applications. The antimicrobial combinations effects are summarized in Table 1.

**Table 1.** Antimicrobial combination effects between Product A and Product B.

| Product A | Product B | Well diffusion assay | Checkerboard Assay |
| --- | --- | --- | --- |
| Type R | Manuka honey | Synergy | Synergy |
|  | MGO | Synergy | Synergy |
|  | Cinn HA | No antimicrobial effect | No antimicrobial effect |
|  | Cinn Oil | No antimicrobial effect | No antimicrobial effect |
|  | Neem seed oil | No synergy | Additive |
| **RE** | Manuka honey | Synergy | Synergy |
|  | MGO | Synergy | Synergy |
|  | Cinn HA | No antimicrobial effect | No antimicrobial effect |
|  | Cinn Oil | No antimicrobial effect | No antimicrobial effect |
|  | Neem seed oil | No synergy | Additive |
| **SWP** | Manuka honey | Additive | Additive |
|  | MGO | Additive | Synergy |
|  | Cinn HA | No antimicrobial effect | No antimicrobial effect |
|  | Cinn Oil | No antimicrobial effect | No antimicrobial effect |
|  | Neem seed oil | No synergy | Additive |
| **SWPE** | Manuka honey | Additive | Synergy |
|  | MGO | Additive | Synergy |
|  | Cinn HA | No antimicrobial effect | No antimicrobial effect |
|  | Cinn Oil | No antimicrobial effect | No antimicrobial effect |
|  | Neem seed oil | No synergy | Additive |
| **Neem seed oil** | MH | No synergy | Antagonistic |
|  | MGO | No synergy | Antagonistic |

It is well-known that dietary sugars contribute to the progression of dental caries and oral biofilm formation (22). Manuka honey, like most honeys has a high sugar content. Although manuka honey was synergistic with cranberry in this study, by substituting Manuka honey with one of its main bioactive agents, MGO, we can avoid sugars, if this is to be used orally as a dentifrice.

One of the main drawbacks of the well diffusion assay is that some natural products with larger molecule sizes could not diffuse into the BHI agar leading to false negative results. The color of cranberry extracts surrounding the well, indicated the level of diffusion into the agar. Cranberry extracts have proanthocyanidins (PAC) that polymerize over time while maintaining their antimicrobial properties. But the polymerized components could not diffuse into the agar in the well diffusion assay. To test synergy between these products, the checkerboard assay was used.

Cranberry PAC that polymerizes to form larger molecules were more bioactive in the checkerboard assay than the well diffusion assay. The larger polymers could not diffuse freely though the agar although, the MIC of these extract were lower than that of the other cranberry extracts. Another important feature was that this PAC-rich extract was antimicrobial even after polymerization. While neem did not show synergy in the well diffusion assay, with small zones of inhibition, it showed more antimicrobial activity in the checkerboard assay. Therefore, while testing natural products it is important to keep in mind the structure of the bioactive molecules with respect to the assay that’s being used to test it.

The checkerboard assay has many advantages. The liquid broth medium avoids the agar diffusion problem faced by products with larger molecule sizes. We can simultaneously test combinations of various concentrations of both the test substances. The MIC and FIC for each antimicrobial combination can be determined in a single plate. It is relatively easy to calculate the FIC index.

Some research (17) suggest that synergy is present when the mean FIC index is ≤ 0.5, additive/ indifference is present when FIC index is between 1 and 4, and antagonism is more than 4. The 0.5 conservative number will account for random effects in a microdilution checkerboard assay. But as Meletiadis et al. describes, a 0.5 FIC is not a natural cut off, making it difficult to classify FICs between 0.5 and 1(21). Experimentally, an FIC index between 0.5 and 1 shows statistically significant synergy (23) and run-to-run values do not vary to a point where synergy is demonstrated in one run and antagonism is seen in another run (24, 25). When the concentrations of a cranberry extract were shifted in this study to find intermediate FIC index values, the mean FIC index remained the same and the lowest FIC index value changed from 0.27 to 0.28. Therefore, the perception that checkerboard data may be more variable that single-antimicrobial data is questionable. So, it seems better to go by the cut-offs indicated in Table 2. To make the study more robust, two other methods of assessing checkerboard assays were also used.

**Table 2.**
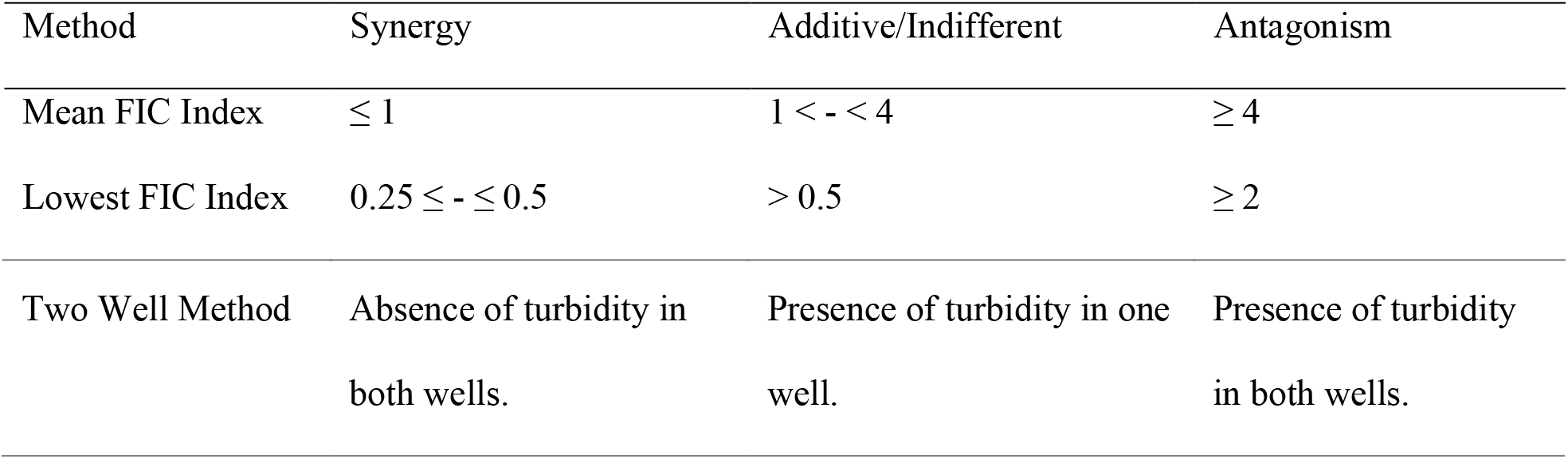
Interpreting the Checkerboard Assay.

**Table 3.**
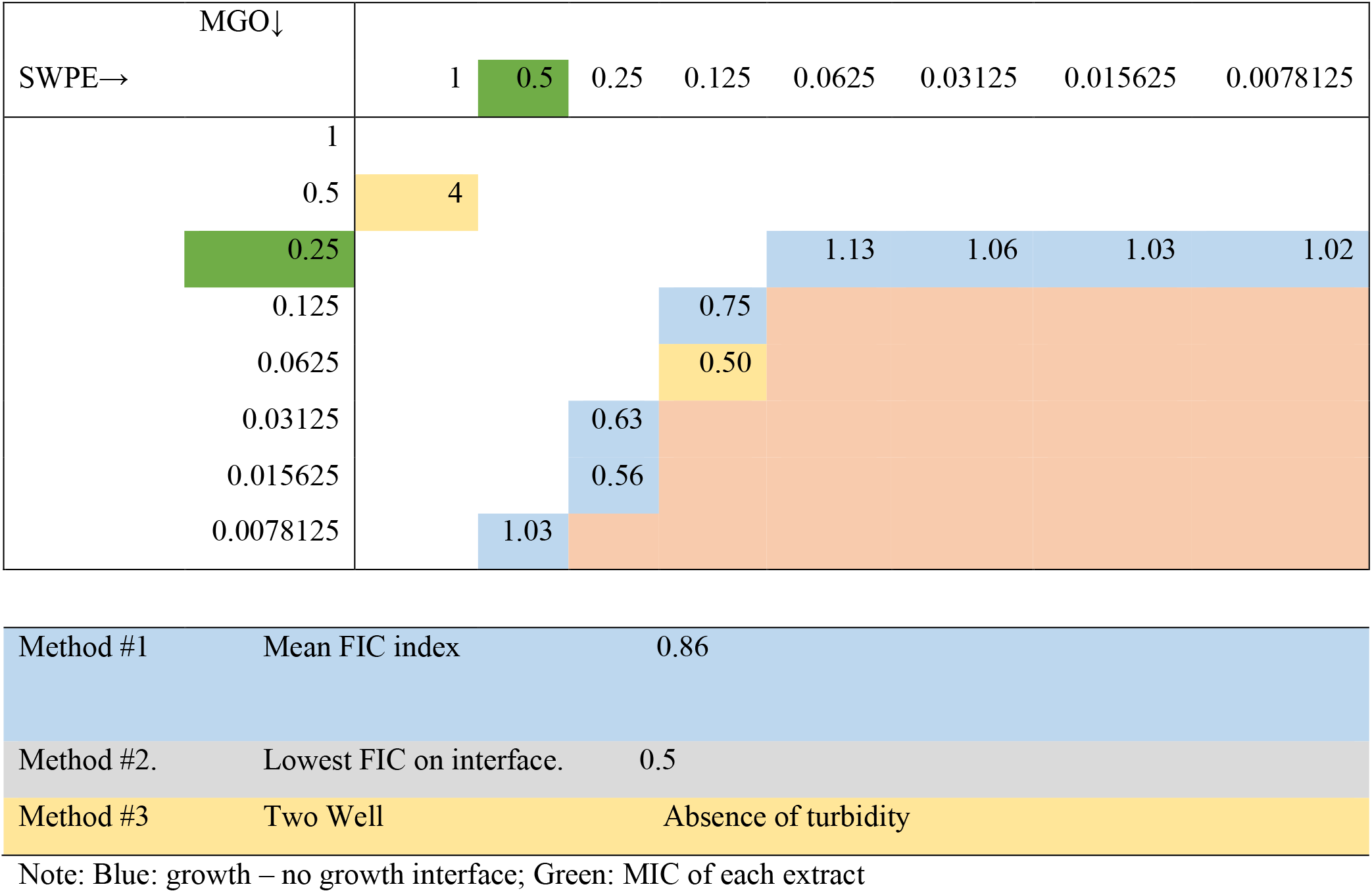
Interpreting the Checkerboard assay with Cranberry extract (SWPE) and MGO

While these results show potential for some synergistic pairs, more research is required before these extracts can be assessed for use as a dentifrice for human patients. The oral biofilm is complex, with multiple species of bacteria embedded in a polymer matrix. The effect of these natural agents on the biofilm should also be studied. A crystal violet biofilm assay can be used to determine the antimicrobial activity of synergistic pairs against the formation of new biofilm. A potential difficulty for using this method to test cranberries or similar compounds, is that the agents have absorption wavelengths close to that of crystal violet and that may lead to difficulties while using a spectrophotometric plate reader. Also, randomized control studies could be performed on small groups of individuals to check the effectiveness of synergistic pairs *in vivo*. The compounds could be used as a mouthwash or incorporated in a toothpaste and tested over time to detect the effects on dental plaque and calculus formation. This would also give more data regarding the effects of saliva on the synergistic combinations. By growing periodontal bacteria anaerobically, and subjecting them to well diffusion and checkerboard assays, we may also find synergistic compounds that can treat gum disease. Other plant derived antimicrobial agents can be tested in both the well diffusion assay and the checkerboard assay. By identifying natural extracts that have proven antimicrobial activity *in vitro* or *in vivo* and combining them with other agents, we can test for synergy. It is important to keep in mind that the likelihood of detecting synergy is higher if the two agents being combined have different antimicrobial mechanisms. If they have similar modes of action, they tend to act in an additive manner and may even be antagonistic.

With the invention of newer technologies, it is crucial to find the right tools that can further the study of antimicrobial synergy and use natural products more effectively to fight tooth decay.

## Acknowledgements

Thanks to Dr. Leland Russell, Professor of Biology, Wichita State University, for guidance about statistical analysis. Thanks to Ms. Fawn Beckman, Microbiological Lab Coordinator, Wichita State University, for expert technical assistance in culturing techniques.

## Supporting Information

